# Combined Effects of Standing and Passive Heating on Attenuating Sitting-Induced Vascular Dysfunction

**DOI:** 10.1101/2020.04.30.060087

**Authors:** Aaron R. Caldwell, Lisa T. Jansen, Megan E. Rosa-Caldwell, Erin K. Howie, Kaitlin M. Gallagher, Ronna C. Turner, Matthew S. Ganio

## Abstract

Recent evidence suggests prolonged sitting strains the arteries through angulations that create turbulent blood flow. This turbulent flow reduces flow-mediated dilation (FMD), a key indicator of vascular health. The purpose of this study was to determine if arterial angulations (through sitting or standing), shear stress (through local heating), or a combination of these factors affected vascular function. In addition, we sought examined the impact of biological sex on these responses.

**Methods:** Twenty-six healthy, young (18-40 years old) males (n=13) and females (n=13) participated. Participants completed two experimental trials (2-h sitting and 2-h standing), and, in a randomized fashion, one leg was passively heated. Flow-mediated dilation (FMD) at the superficial femoral artery (SFA), and central and peripheral pulse-wave velocity (PWV) were measured using vascular ultrasound.

**Results:** There was a non-significant decrease in FMD (- 1.48%; *p =* .06) during sitting and the decline in FMD was not different between biological sexes (−1.96% vs -0.93%; *p =* .49, males and females respectively). Passive heating (1.42%; *p* < .05) and standing (1.42%; *p* < .05) both improved FMD in comparison to sitting. However, standing resulted in a significant increase in peripheral PWV (50 cm/s; *p* < .05). Interestingly, the standing was not well tolerated among female participants with seven participants having to stop their initial trial early due to lightheadedness.

**Conclusions:** Both interventions appear to be equally effective at mitigating reductions FMD associated with sitting, but standing increased peripheral PWV. In addition, it does not appear that biological sex moderates these physiological responses.

**New Findings:** *What is the central question of this study?:* Prolonged sitting can cause acutely vascular dysfunction while interventions such as local heating or standing have been explored they have not been used in combination and the role of biological sex has not been fully explored.

*What is the main finding and its importance?:* In this study we demonstrate that either local heating or standing are effective at reducing some of the vascular dysfunction associated with prolonged sitting. Biological sex did not appear appear to play a role in this response. However, standing may also cause some negative effects such as increased arterial stiffness and increase the risk of syncope.

## Introduction

Prolonged periods of sitting (>1 hour) can directly cause acute vascular dysfunction (Thosar et al., 2015). Additionally, when physically active adults are restricted to sitting for five days, significant vascular dysfunction begins to manifest suggesting more chronic effects (McManus et al., 2015; Teixeira et al., 2017). While there is a clear negative impact of sitting, the specific mechanisms involved in the development of this “proatherogenic hemodynamic environment” (Padilla & Fadel, 2017) are largely unknown, thereby making it difficult to develop strategies to counteract sitting induced vascular dysfunction.

Specific vascular-related consequences can occur in just a few hours of inactivity. In 1 to 4 hours, sitting reduces vascular function by up to 4%, as measured by flow mediated dilation (FMD) (Restaino et al., 2015; Thosar et al., 2015). This reduced vascular function may be partially moderated by a systemic vasoconstrictor response, as evidenced by increased endothelin-1(ET-1) after 2 hours of sitting (Ballard et al., 2017). In addition, arterial stiffness, as measured by pulse wave velocity (PWV), increases in the lower limbs after 2 hours of standing (Caldwell et al., 2018).

Current evidence suggest that the two primary factors that modify vascular function during prolonged sitting are shear rate (i.e., directional force of blood flow along the vessel wall) and arterial angulation (i.e., bending of the artery due to limb position) (Padilla & Fadel, 2017). FMD, as a measure of vascular function in particular seems to be sensitive to both factors(Morishima et al., 2017; Restaino et al., 2016). However, lower limb arterial stiffness (i.e., PWV) appears to increase even in the absence of arterial bending (i.e., during standing) (Caldwell et al., 2018). Therefore, standing may not always be beneficial and, in fact, may have some negative effects on the vasculature

Standing workstations have been promoted as a healthier alternative to sitting (Lopez-Jimenez, 2015), because standing may attenuate sitting-induced reductions in FMD(Morishima et al., 2017). Reduction in arterial angulation that occur during standing may be the primary mechanism underlying. Generally, improved vascular function compared to standing. he hip and knee joints are bent at 90° in the seated position. Leg blood flow is severely reduced when the legs are bent at the knee compared to when individuals are fully supine (0° angle) (Morishima et al., 2017). This would suggest that proximal arterial bending reduces lower limb blood flow and could explain much of the vascular dysfunction that occurs after sitting. Elimination of these arterial angulations, by standing, may reduce turbulent blood flow in the femoral artery compared with sitting (Walsh et al., 2017).

Additionally, prolonged standing (Morishima et al., 2017), as well as prolonged sitting (Restaino et al., 2015), reduce shear rate in the femoral artery. When local heating is applied during prolonged siting, FMD increases because antegrade (forward moving) shear rate is increased and retrograde (backward moving) shear rate is reduced (Restaino et al., 2016; Teixeira et al., 2017). However, it is unknown if increasing shear rate during prolonged standing affects vascular function. It is possible that local heating with standing could counteract possible detrimental side effects of standing (e.g., increased arterial stiffness).

Biological sex is known to strongly influence vascular function, but very few studies to date have considered biological sex as a potential variable of interest when evaluating sitting induced vascular function (McManus et al., 2015; Vranish et al., 2017). While standing and local heating have been promoted as potential interventions or alternatives to protect against sitting-induced vascular dysfunction, these studies have primarily used males as research participants (Morishima et al., 2017; Restaino et al., 2016). Therefore, it is unknown if increasing shear rate through local heating, or altering shear rate, via prolonged standing, affect males and females differently. The limited research available suggests that estrogen may play a role since prepubertal girls are not protected from sitting induced vascular dysfunction (McManus et al., 2015). In the studies that have included female participants (n=6 studies), only one has examined potential differences between the sexes (Vranish et al., 2017). This is particularly worrisome considering female participants are partially protected from sitting induced vascular dysfunction(Vranish et al., 2017). Considering vascular dysfunction manifests differently in male and female participants, it follows that biological sex may interact with the interventions such as standing or local heating. There is emerging evidence that biological sex influences vascular function following prolonged sitting. However, it is unknown to what degree sitting induced vascular dysfunction, or interventions to prevent it (e.g., standing or local heating), are affected by biological sex.

Currently, there is a gap in the literature documenting the specific effects of prolonged standing on both local (FMD and PWV) and systemic vascular function (ET-1). Therefore, the purpose of this study is to determine the local (PWV and FMD) and systemic (ET-1) vascular responses to both sitting and standing. We hypothesized that standing, compared to sitting, would attenuate reductions in FMD compared to sitting (i.e., interaction between time and trial) due to the removal of the arterial angulations that in turn reduce retrograde shear rate. However, we also hypothesized that PWV and ET-1 responses would be similar between sitting and standing due to increased vasoconstriction. Lastly, we hypothesized that local heating would increase shear rate and lead to improvements in FMD and PWV compared to sitting or standing without local heating. A secondary aim of this study was to investigate if biological sex moderates the vascular responses to interventions such as standing and heating. We hypothesized that, similar to previous studies, females would have attenuated vascular dysfunction in response to 2 hours of sitting relative to males. In addition, we estimated the interaction between biological sex and interventions (i.e., local heating and standing) intended to prevent sitting induced vascular dysfunction (i.e., Sex x Intervention interaction). In light of the available evidence, we hypothesized that males would have a greater improvement in FMD during standing while there would be equal improvements in FMD between sexes from local heating.

## Methods

Non-obese males and females between the ages of 18-40 years old were recruited to participate in this study. Exclusionary criteria included previous history of cardiovascular issues, hypertension, current drug use that can affect endothelial function (i.e., SSRIs, β-blockers, nitrates, and calcium channel blockers), obesity (BMI>30 kg/m^2^), smoking, or a history of problems with fainting or dizziness(Harris et al., 2010; Thijssen et al., 2011). To, control for the effects of the menstrual cycle, female participants were only included if they were taking regular monophasic hormonal contraception (Harris et al., 2010; Thijssen et al., 2011). A description of the sample can be found within Table 1.

**Table 1.** Participant Characteristics (Mean ± SD)

| **Table 1. Participant Characteristics (Mean ± SD)** |  |  |
| --- | --- | --- |
|  | Female | Male |
| Age (years) | 26 ± 3 | 24 ± 3 |
| Body Mass Index (kg/m<sup>2</sup>) | 21.1 ± 2.3 | 25 ± 3.2* |
| Body Fat<sup>a</sup> (%) | 27.0 ± 5.9 | 21.9 ± 7.3 |
| Physical Activity<br>(MET·min/week) | 2696 ± 3364 | 2565 ± 1767 |

<sup>a</sup>Obtained via DXA scan
<sup>b</sup>Obtained from International Physical Activity Questionnaire
\*Significant difference from females ( $p < .05$ )

### Experimental Protocol

All activities were approved by the University of Arkansas Institutional Review Board(). At a familiarization visit, participants signed an informed consent that approved by the University of Arkansas Institutional Review Board, and completed a medical history questionnaire. In accordance with current guidelines for FMD (Harris et al., 2010; Thijssen et al., 2011), participants were asked to refrain from alcohol and exercise for 8 hours, and food for 6 hours prior to each experimental trial. Pre-test compliance was verified with a 24-hour history questionnaire.

A schematic of the protocol design is in Figure 1. Each participant completed two experimental trials (completed in random counterbalanced order): sitting (2 hours of sitting), standing (2 hours of standing). Moreover, in each trial one leg was passively heated while another acted as a control. This methodology provides a time control and is often used in this area of research (Morishima et al., 2017; Restaino et al., 2015, 2016; Thosar et al., 2015; Walsh et al., 2017). Further, previous literature has demonstrated 2 hours is sufficient to induce observable vascular dysfunction during sitting (Thosar et al., 2015), and internal pilot data has indicated that standing in the same place for > 2 hours is not tolerable for many individuals. In the standing trial, participants stood while watching television while in the sitting trial participants remained in a standard phlebotomy chair while watching television.

**Figure 1.**
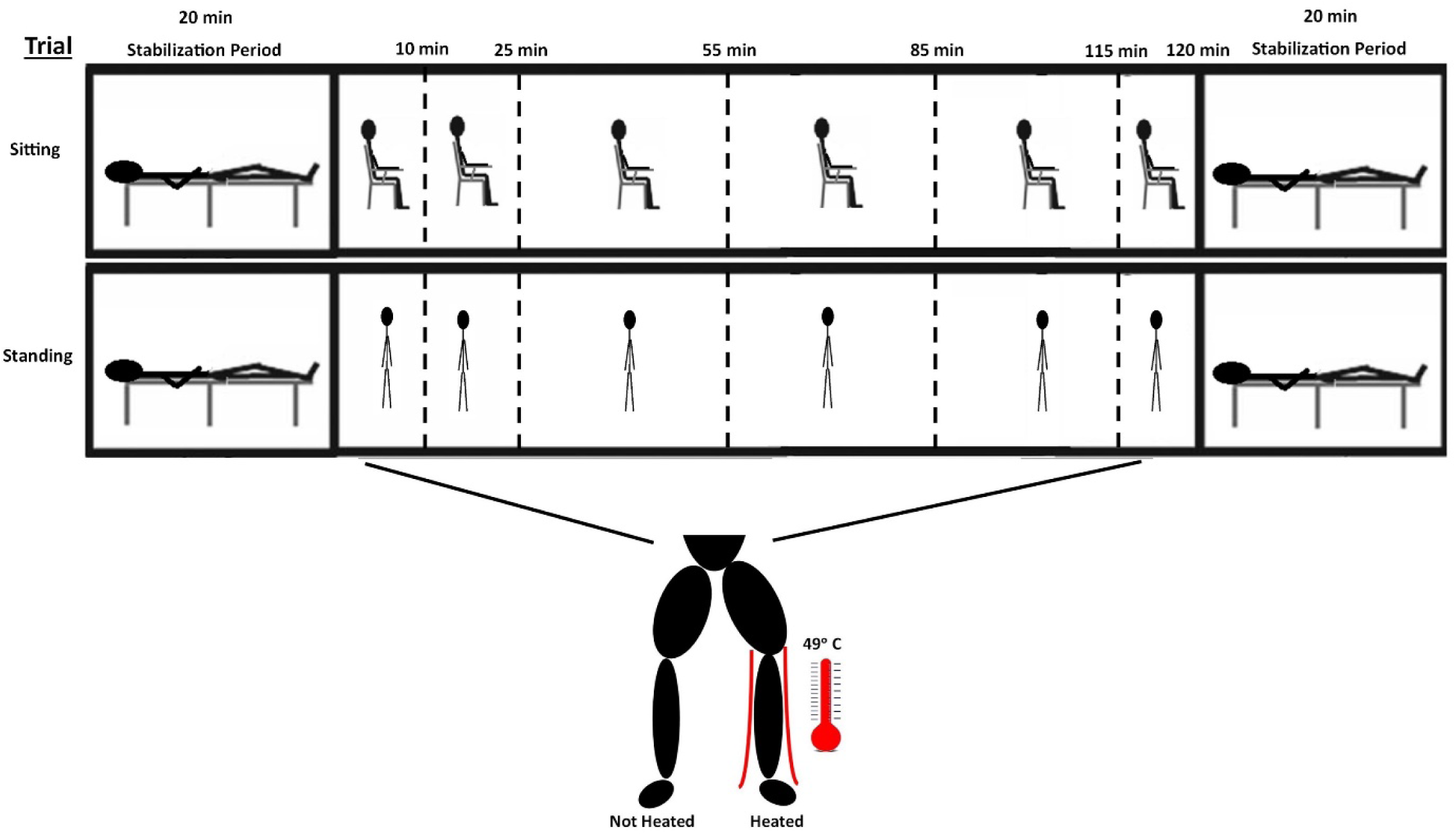
Study Design. Dashed lines represent the measurement of shear rate and blood flow at the superficial femoral artery (both legs) over the course of 2 minutes. During each trial, one leg was heated via a water-perfused pant leg circulating 49°C water. Trials (sitting vs standing) and passive heating (left leg vs right leg) was completed in a randomized, counter-balanced order.

#### Local heating

In order to passively heat the legs, participants were dressed in a water-perfused, tube-lined pants that covers the intervention leg (Allen-Vanguard, Ottawa, Canada). The suit permits control of skin temperature by changing the temperature of the water perfusing the suit. The local heating leg, after baseline measures, had hot water (49°C) perfuse the pants throughout the standing or sitting bout. Skin temperature was recorded on both legs at the thigh (midway between the patella and femoral head), lateral calf (midway between the lateral malleolus and head of the fibula), and on the anterior of the foot using six (three on each leg), small, wireless thermichrons that recorded skin temperature (iButtons, Maxim Integrated, San Jose, CA).

Immediately before and after each trial, participants laid supine, on a padded phlebotomy chair and rested for 20 minutes prior to the measurement of baseline FMD, shear rate, blood pressure and heart rate. This rest period provided subject stabilization prior to FMD^14^. Ultrasound imaging (for shear rate measures) at the superficial femoral artery (SFA) and other cardiovascular measures (described below) were measured at minutes 10, 25, 55, and 115 of each trial. Blood pressure and heart rate measurements were obtained in duplicate during measurements of shear rate and FMD.

### Experimental Measures

#### Pulse wave velocity

Arterial stiffness was indexed using PWV, which is the preferred method^17^. Central PWV was calculated from the carotid and femoral arteries, while Peripheral PWV of the lower limb was calculated from femoral and dorsalis pedis arteries. PWV was measured with Doppler ultrasound (GE GoldSeal LOGIQ eBT08) using the foot-to-foot method (Calabia et al., 2011). Specifically, PWV was calculated as the distance between measurement sites divided by the time delay between the two waveforms. The distance between arterial sites for PWV was calculated as the direct distance between the femoral and dorsalis pedis sites. Distances between sites was calculated for each individual trial by measuring from the distal edge of the ultrasound probe with a retractable cloth tape measure. All PWV measures were performed on both sides of the body (left and right) in each trial with consistent probe location being assured by marking the skin with a surgical marker. A three-lead (RL, LA, and RA leads) ECG was utilized to calculate the time delay from the R-wave to the foot of the pulse wave.

In this study, upright (sitting and standing) measures of PWV were obtained. Body posture influences hemodynamic regulation of parameters related to arterial stiffness (Reesink et al., 2007), but measuring arterial stiffness in the upright position is considered reliable (Nürnberger et al., 2011). Therefore, upright measures were only compared to other upright measures. Measurement site order was randomized between participants but consistent within-participant for each trial.

#### Flow-mediated dilation and shear rate measurements

All FMD and shear rate measurements were obtained at the SFA on both legs. Artery diameter and blood velocity was measured using duplex-Doppler ultrasound. Simultaneous diameter and velocity signals were obtained in duplex (Thijssen et al., 2011). During the standing or sitting bouts, ultrasound videos to measure shear rate and artery diameter at the SFA were recorded for one minute. These ultrasound images were analyzed with antegrade (forward moving) and retrograde (backward moving) shear rate being recorded as well as artery diameter. Prior to FMD measurement, an automatic rapid inflation cuff (E-20 rapid cuff inflator; D.E. Hokanson, Bellevue, WA) was placed on their right thigh directly distal to the knee joint, and, therefore, distal to the SFA recording location. After baseline FMD was assessed, the placement of the ultrasound probe was marked with a surgical pen to ensure consistency of subsequent measurements. Two minutes of baseline hemodynamics was recorded, the cuff was then inflated to a pressure of 250 mmHg for 5 minutes, and the vasodilatory response was then recorded for 5 minutes after cuff release. A validated(Faita et al., 2011) software (FMD Studio, Quipu, Pisa, Italy) for beat-to-beat analysis vessel wall detection and quantification of shear rate was utilized. Shear rate was calculated from the blood flow velocity and frame-by-frame vessel diameter from the duplex ultrasound, as previously reported(Thijssen et al., 2011). Furthermore, in order to account for allometric scaling, the slope between baseline diameter and maximal diameter was assessed (Atkinson & Batterham, 2013a). Overall, the slope between these diameters was 0.89 and this slope was consistent between groups and time points. Therefore, FMD was calculated as 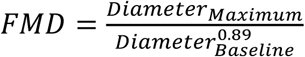 consistent with previously reported adjustments for allometeric scaling (Atkinson & Batterham, 2013b).

#### Other Cardiovascular Measures

Participants were fitted with an automated blood pressure cuff (Tango+; SunTech Medical, Inc., Morrisville, NC, USA) placed on the right brachial artery to obtain blood pressure via electrosphygmomanometry. Mean arterial pressure (MAP) was calculated as two-thirds diastolic pressure plus one-third systolic pressure and averaged from duplicate blood pressure measurements taken at each time point. Blood pressure and HR measurements were measured in duplicate during measurements of shear rate and FMD.

After the 20 minute rest period and FMD measurements, blood was collected from the antecubital vein through venipuncture. Blood was allowed to clot for at least 15 minutes, after which blood was centrifuged at 3000G and serum was collect for measurement of ET-1. ET-1 was quantified using a commercial kit (ThermoFisher, Cat#EIAET1) and standard plate reading instruments (HTX Multi-Mode Reader, Biotek Synergy) and software (Gen5, Biotek Synergy).

### Statistical Analysis

To compare effect of biological sex on FMD response to sitting, a linear mixed model with time and biological sex as factors was utilized (time x sex) while only including data from the sitting, non-heated leg. Each specific aim of this study was analyzed using a linear mixed models to analyze differences in shear rate, vessel diameter, and FMD between trials (i.e. condition: sitting vs. standing, or intervention: heating vs. no heating) and subjects (i.e., sex) while controlling for baseline values (i.e., baseline time varying covariate) (Winer et al., 1991). When a significant main effect or interaction was observed, the estimated marginal means were then inspected for significant pairwise comparisons. When significant effects were observed, differences are reported as the mean difference with 95% confidence interval (95% C.I.). In addition, descriptive statistics (e.g., BMI, age, and body fat percentage) between males and females were compared using an independent samples t-test. Significance was set at *p* < .05.

## Results

### Hemodynamics during sitting vs standing and heating vs non-heated

There was no effect of biological sex on MAP in response to sitting or standing. However, MAP was elevated from the supine measures (baseline and post) during standing or sitting (8.4 mmHg 95% C.I. [6.8, 9.9], *p* < .001), and standing elevated MAP compared to sitting (4.9 mmHg 95% C.I. [2.2, 7.7], *p* < .001). HR was elevated when participants were upright compared to the supine measures (19 bpm 95% C.I. [17, 22]; *p* < .001), and during the standing trial compared to sitting (6 bpm 95% C.I. [2, 9]; *p =*.004) at all upright time points. However, the increase in HR was moderated by biological sex (interaction between Sex x Trial; *p =*.002). Females had significantly higher HR during the standing trial compared to males (11 bpm 95% C.I. [1, 22]; *p =*.049), but there were no differences between sexes during the sitting trial (2 bpm 95% C.I. [-9, 13]; *p =* .69). We also note that the standing trial was not well tolerated. A total of 8 participants reported lightheadedness causing the trial to be cancelled and rescheduled. However, they seemed to tolerate the experimental trial on the 2^nd^ attempt. Most of the participants that reported lightheadedness (7/8) were females.

During sitting and standing, passive heating increased antegrade shear rate (72.8 s^-1^ 95% C.I. [58.9, 86.6]; *p* < .001) to a similar extent in both male and female participants. This increase was sustained throughout the trials (i.e., main effect of intervention) while there were no differences between sitting and standing (i.e. no main effect of trial). Retrograde shear rate was reduced during passive heating compared to the control limb (13.2 s^-1^ 95% C.I. [10.7, 15.7]; *p* < .001) and during standing compared to sitting (3.1 s^-1^ 95% C.I. [0.23, 5.95]; *p =* .035). In contrast, superficial femoral artery diameter was reduced during standing compared to sitting (−0.28 mm 95% C.I. [-0.48, -0.08]; *p =* .005), but there was no effect of passive heating or biological sex.

### Effectiveness of Local Heating during the Experimental Trials

Skin temperature at the calf increased at the initiation of passive heating, and peaked at 40.1 °C, 95% C.I. [39.9, 40.3]. Similarly, skin foot temperature increased during passive heating and peaked at 35.0 °C, 95% C.I. [34.5, 35.5]. While thigh skin temperature did not increase throughout the trial (i.e., no effect of time; *p* > .05); thigh temperature on the heated leg (31.9 °C, 95% C.I. [31.6, 32.3]) was greater than the control (31.3 °C, 95% C.I. [30.9, 31.9]; main effect of intervention: 0.63 °C 95% C.I. [0.63, 0.72]; *p* < .001). Changes in skin temperature at all sites (thigh, calf, and foot) were not different between biological sexes (*p* > .05).

### Pulse Wave Velocity

Overall, peripheral PWV was higher in the standing trial whether in the supine (48 cm/s 95% C.I. [15, 80]; *p =* .004) or upright position (76 cm/s 95% C.I. [25, 134]; *p =* .007). However, there was no effect of passive heating (*p* > .05) or biological sex (*p* > .05) (Figures 2C and 2D). As for central PWV, there was no significant effect of trial (sitting vs standing), intervention (passive heating), or biological sex (Figures 2A and 2B).

**Figure 2.**
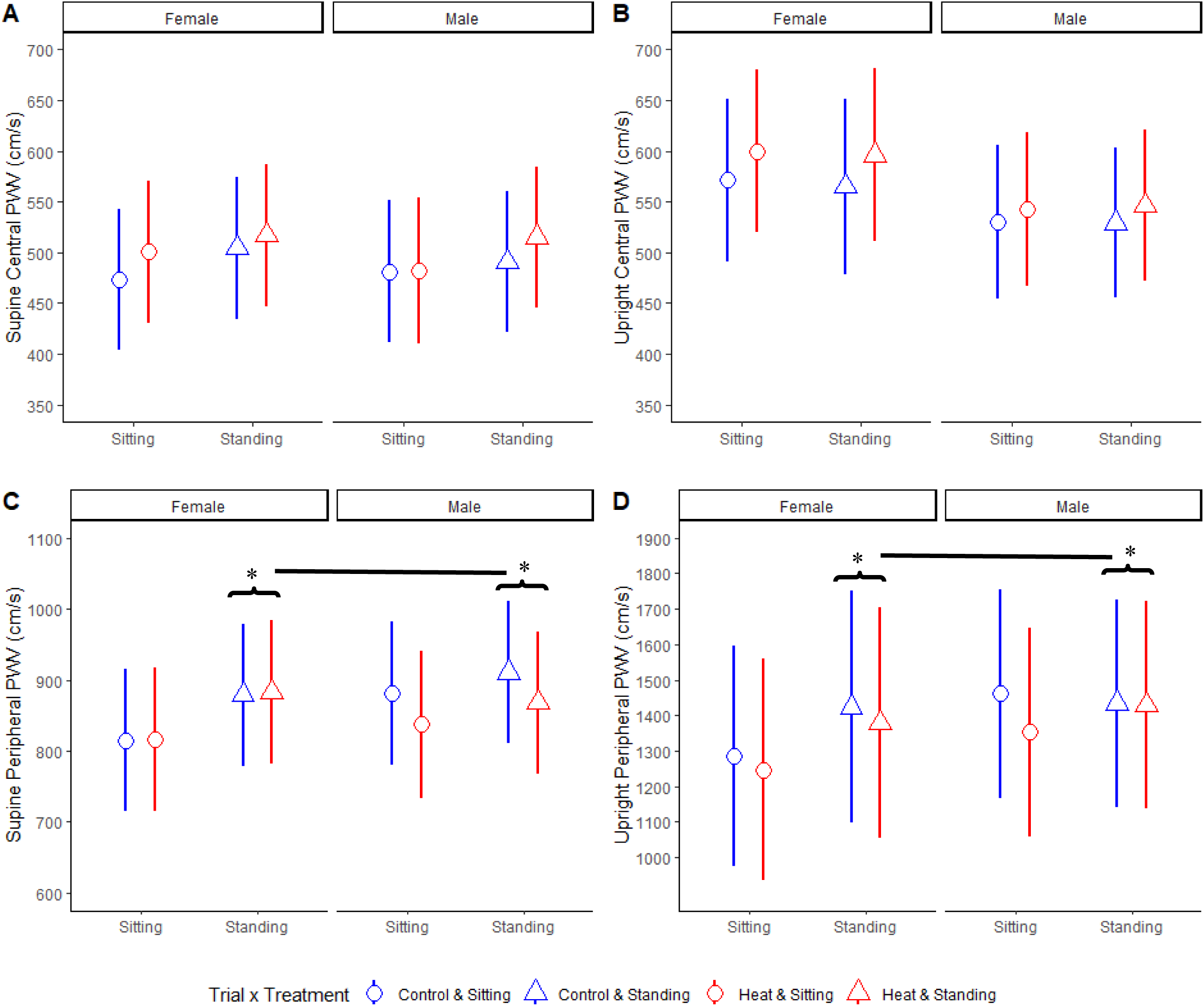
Measures of arterial stiffness. Central PWV while (A) supine and (B) upright and peripheral PWV while (C) supine and (D) upright. *Indicates a significant difference from sitting (*p* < .05). All data presented as (mean ± SD).

### Flow-Mediated Dilation

#### Influence of biological sex on responses to sitting

Overall, there was a non-significant decrease in FMD in the sitting trial (−1.45% 95% C.I. [-2.95, .05]; *p =* .061). Further, there was no effect of biological sex on the FMD response to sitting (Table 3). Baseline artery diameter (i.e., prior to ischemia) significantly increased in the sitting trial (0.15 mm 95% C.I. [0.01, .28]; *p =* .032) and males tended to have a larger SFA diameter (1.01 mm 95% C.I. [0.49, 1.53]; *p* < .001); independent of the time point (i.e., main effect of sex; Table 3). Further, maximum shear rate during FMD tended to be greater in the female participants (190 s^-1^ 95% C.I. [75, 306]; *p =* .002; Table 3).

**Table 2.**
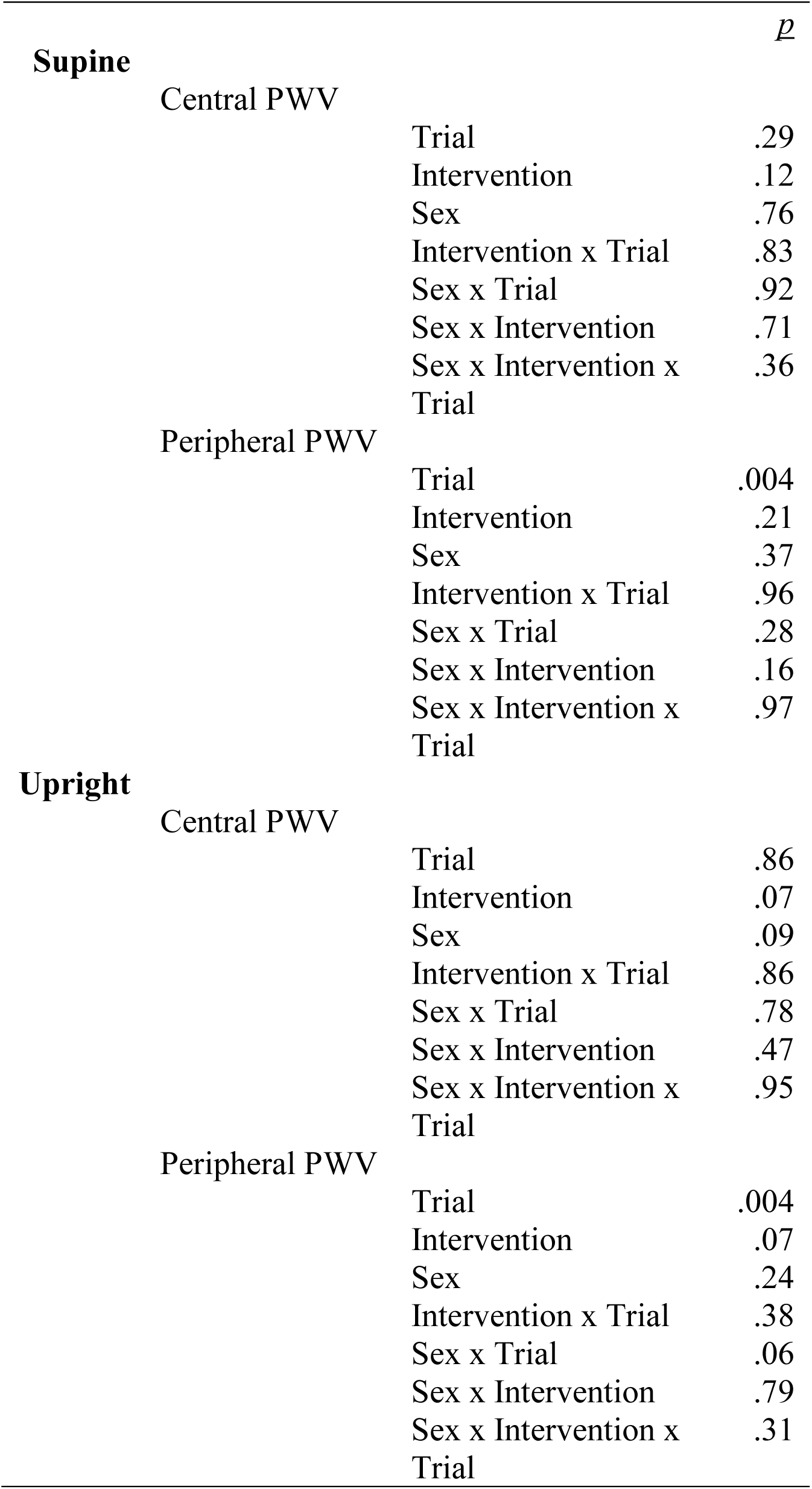
Combined effects of sitting, standing, and biological sex: Pulse wave velocity linear mixed model ANOVAs

**Table 3.** Influence of biological sex on responses to sitting: Flow-mediated dilation linear mixed model ANOVAs

|  |  | *p* |
| --- | --- | --- |
| FMD |  |  |
|  | Sex | .65 |
|  | Time | .06 |
|  | Time x Sex | .49 |
| Baseline Diameter |  |  |
|  | Sex | <.001 |
|  | Time | .031 |
|  | Time x Sex | .38 |
| Maximum Shear Rate |  |  |
|  | Sex | <.001 |
|  | Time | .19 |
|  | Time x Sex | .99 |

#### Influence of biological sex on responses to passive heating and standing

Overall, there was a main effect of both trial and intervention (Table 4; both *p* < .05). Passive heating resulted in higher FMD compared to control (1.64% 95% C.I. [0.39, 2.89]; *p =* .011; Figure 3A), independent of trial (sitting and standing). Further, standing resulted in higher FMD (1.42% 95% C.I. [0.18, 2.66]; *p =* .025; Figure 3A). Male and female participants responded similarly to the passive heating and standing (i.e., no interactions between sex and other factors; Table 4). Further, baseline diameter was lower after the standing trial, independent of intervention and biological sex (0.23 mm 95% C.I. [0.10, 0.36]; *p* < .001; Figure 3B). In addition, maximum shear rate during FMD was greater in the passively heated limb in the female participants (320 s^-1^ 95% C.I. [139, 502]; *p* < .001; Figure 3C), but there was no significant change in the male participants (35 s^-1^ 95% C.I. [-144, 213]; *p =* .69; Figure 3C).

**Table 4.** Combined effects of biological sex, heating, and standing: Flow-mediated dilation linear mixed model ANOVAs

|  |  | *p* |
| --- | --- | --- |
| FMD | Sex | .99 |
|  | Trial | .024 |
|  | Intervention | .011 |
|  | Sex x Trial | .97 |
|  | Sex x Intervention | .75 |
|  | Trial x Intervention | .39 |
|  | Sex x Trial x Intervention | .18 |
| Baseline Diameter | Sex | .34 |
|  | Trial | <.001 |
|  | Intervention | .15 |
|  | Sex x Trial | .13 |
|  | Sex x Intervention | .27 |
|  | Trial x Intervention | .33 |
|  | Sex x Trial x Intervention | .70 |
| Maximum Shear Rate | Sex | .003 |
|  | Trial | .10 |
|  | Intervention | .007 |
|  | Sex x Trial | .39 |
|  | Sex x Intervention | .028 |
|  | Trial x Intervention | .29 |
|  | Sex x Trial x Intervention | .40 |

**Figure 3.**
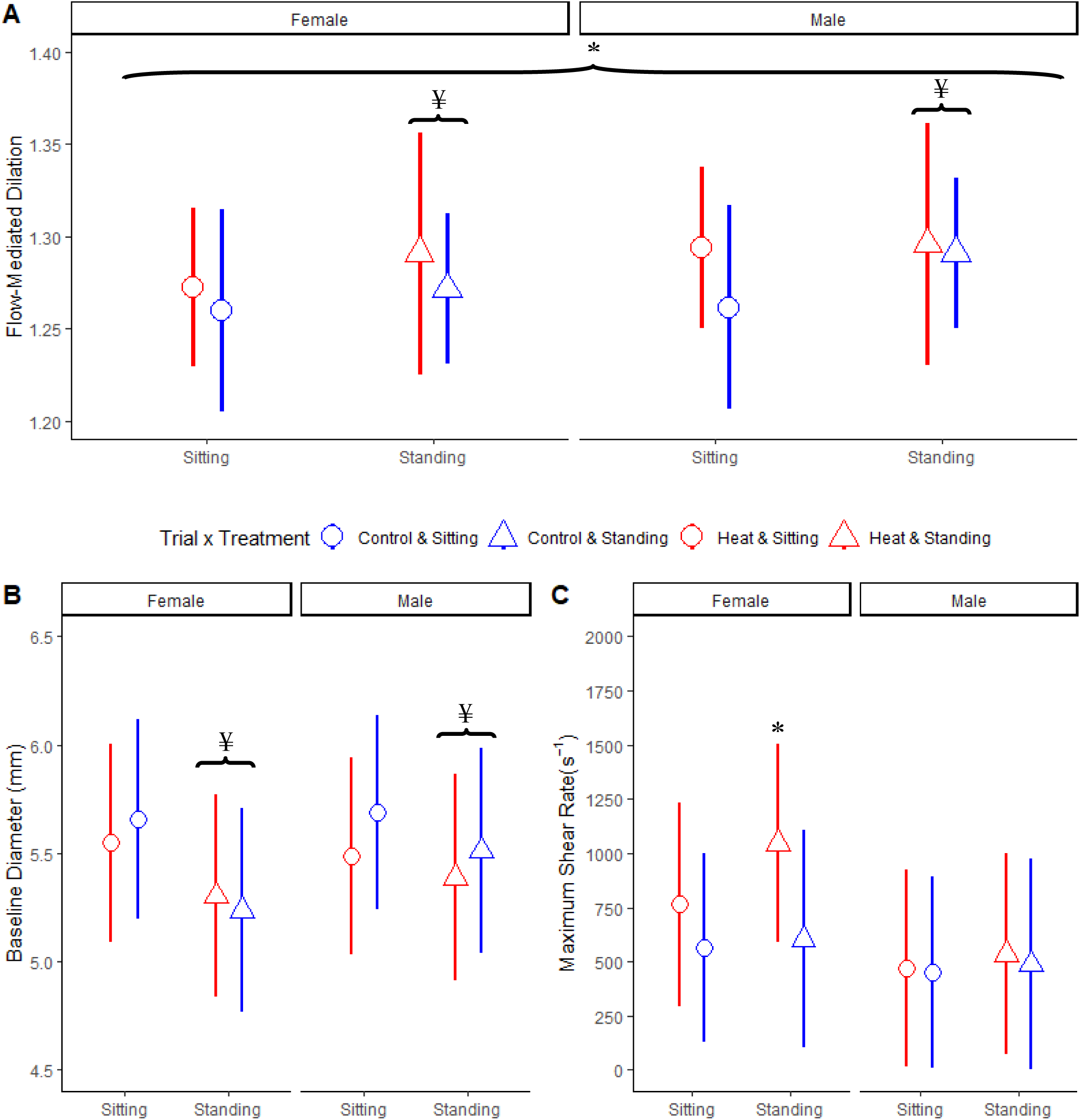
(A) Flow-mediated dilation (B) Baseline diameter and (C) maximum shear rate at post-trial (after 2-h sitting/standing) after controlling for baseline differences (i.e., adjusting for baseline as a covariate). *Indicates a significant difference from control (*p* < .05) and ^¥^ indicates a significant difference from sitting (*p* < .05). All data presented as (mean ± SD).

### Endothelin-1

Overall, there were no significant changes in ET-1 over the course of 2 hours of sitting (−0.56 pg/mL 95% C.I. [-2.26, 1.13]; *p =* .45), or standing (−0.63 pg/mL 95% C.I. [-2.33, 1.05]; *p =* .51). There were no differences between the males and females (0.74 pg/mL 95% C.I. [-1.39, 2.87]; *p =* .47), after accounting for baseline differences, in ET-1 (Figure 4).

**Figure 4.**
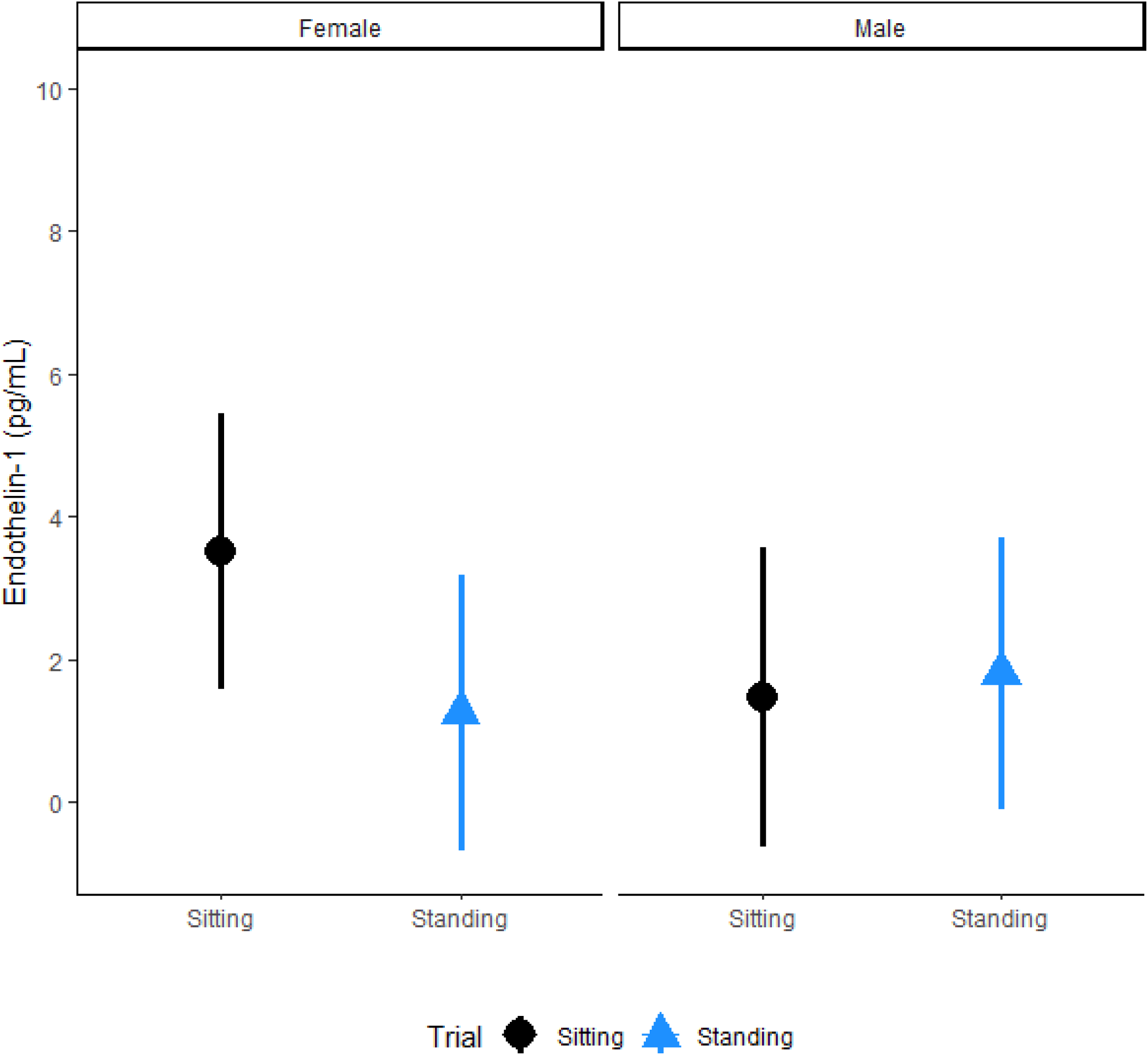
Changes in ET-1 (mean ± SD) after 2 hours of sitting and standing in male and female participants while controlling for baseline differences in ET-1 (baseline as a covariate).

## Discussion

The primary finding of the present study is that leg endothelial function, as assessed with FMD, is improved relative to sitting when healthy, young adults, with normal resting FMD, stand or have the lower limbs passively heated. The combination of passive heating and standing did not confer a significant additional benefit compared to standing or passive heating alone. Peripheral PWV, an index of lower limb arterial stiffness, increased during standing but not sitting. Passive heating did not attenuate the increases in peripheral PWV during standing. In addition, in this study we compared the effects of prolonged sitting and interventions (heat and standing) FMD, PWV, and ET-1 between sexes. Contrary to previous work, we found the responses in FMD, PWV, and ET-1 were similar between males and females. Collectively, our findings indicate a lack of sex differences in vascular responses to 2-h of prolonged sitting or standing. Overall, these findings indicate that FMD is improved by modifying shear rate (standing and passive heating), peripheral PWV increases during standing, and biological sex does not moderate these responses.

Previous research indicates females are protected from sitting-related impairments in FMD, despite both male and female participants experiencing similar reductions in shear rate during a 3-h sitting period (Vranish et al., 2017). Instead, our data indicate males and females experience similar reductions in shear rate and FMD during periods of prolonged sitting. However, in our study, we did not induce a significant reduction in FMD after 2-h of sitting. Therefore, the difference between males and females after 2-h of sitting may be masked by the lack of an overall change in FMD during the seated trials.

Shear rate in the femoral artery is reduced during prolonged standing^10^ and sitting^5^. In the current study, when local heating was applied, antegrade shear rate increased and retrograde shear rate was reduced. Interestingly, FMD improved compared to sitting alone in both males and females. Further, retrograde shear rate is reduced during standing, compared to sitting alone. This, in similar fashion to heating, corresponded to an improvement in FMD compared to sitting irrespective of biological sex.

Previous research would indicate that when retrograde shear rate is increased, FMD is reduced (Padilla et al., 2011; Schreuder et al., 2015). *In vivo*, increased retrograde shear rate promotes the development of atherosclerosis (Ziegler et al., 1998). Further, sitting induced vascular dysfunction, as measured by FMD, is eliminated when antegrade shear rate is increased via local heating (Padilla et al., 2011). This supports the view that sitting-induced leg endothelial dysfunction is mediated by a reduction in shear rate in both males and females (Teixeira et al., 2017). Our data lend support to this hypothesis and confirm that local passive heating is effective at increasing antegrade shear rate. Further, standing, while it did not increase antegrade shear rate, did reduce retrograde shear rate in comparison to sitting. The reduction in retrograde shear rate appears sufficient to improve FMD in comparison to sitting. Furthermore, the effects of standing on FMD were similar between males and females. Lastly, the combination of standing and passive heating did not result in differences in shear rate or FMD compared to passive heating alone in males and females (Figure 3).

The current study did not find supporting evidence for the hypothesis that males and females have different vascular responses to prolonged sitting. However, there is evidence from other studies that indicate that females may still be protected from vascular damage. Nitric oxide, a potent vasodilator derived from the endothelium, production and nitric oxide synthase is greater in females than males (Forte et al., 1998). Some have postulated that this greater nitric oxide availability protects females from sitting induced vascular dysfunction despite similar reductions in antegrade shear rate during sitting (Vranish et al., 2017). Further, the sex differences in nitric oxide availability are typically attributed to the increased presence of estrogen in females (Chambliss & Shaul, 2002). As evidence of this, pre-pubertal females have a similar reduction in FMD as adult males (McManus et al., 2015).

Standing also appeared to increase arterial stiffness localized to the lower limbs (peripheral PWV; Figure 2), and this is in agreement with previous studies on the effects of prolonged standing (Caldwell et al., 2018). The increase in peripheral PWV observed during prolonged standing (Figure 2) may be related to the edema that occurred during standing. In fact, we observed greater vasoconstriction (reduction in femoral artery diameter) during the standing trial compared to the sitting trial. However, HR and MAP were elevated during standing compared to sitting (Figure 2). This possibly indicates that during standing venous return is reduced, and total peripheral resistance, as well as HR, are increased in order to maintain cardiac output. The pooling of blood in the lower limbs likely stretched the small veins causing a venoarteriolar response resulting in vasoconstriction(Brothers et al., 2009; Henriksen & Sejrsen, 1976). During vasoconstriction, SFA diameter decreased (Figure 3B) whereas pressure within the femoral artery increased. The increase in pressure, within this specific arterial branch, may have increased peripheral PWV(Nichols & Edwards, 2001). Furthermore, the degree of vasoconstriction was similar between biological sexes, and the corresponding increase in peripheral PWV was similar between males and females. However, there are a myriad of other physiological factors that can affect arterial stiffness (Townsend et al., 2015) that may have also played a role.

The adoption of standing desks is now being promoted as a healthier alternative to sitting (Trinity, 2017). Unfortunately, occupations that require standing are associated with increased low back pain and chronic venous insufficiency (Waters & Dick, 2015). In our study, eight participants were unable to complete the 2-h standing protocol upon their first attempt due to lightheadedness and had to re-schedule this trial for a later date. This effect was likely exacerbated by the participants being fasted and only being allowed minimal movement (shifting weight from one leg to the other). Interestingly, seven of these eight participants were female thus indicating females may not benefit from prolonged periods of standing compared to males. This is not necessarily surprising considering young females are much more likely than young males to experience orthostatic hypotension and syncope (Joyner et al., 2016). In fact, 50% of female medical students, by age 25, report having experienced an episode of orthostatic intolerance (Ganzeboom et al., 2003).

Current evidence suggests an imbalance between α-adrenergic vasoconstriction and β-adrenergic vasodilator is the primary cause of orthostatic intolerance in females (Joyner et al., 2016). In young males, an orthostatic challenge increases sympathetic activity leading to vasoconstriction in order to increase peripheral resistance thereby maintaining MAP despite lower cardiac output (due to reduced venous return). However, due to the differences in adrenergic receptors in young females, the increase in peripheral resistance is blunted and therefore must be offset by increasing cardiac output (as evidenced by an increase in HR). Yet we must acknowledge that we have a relatively limited sample size for this portion of the study as orthostatic intolerance was not our primary outcome variable. As such, future works should seek to specifically quantify the prevalence of this orthostatic intolerance while using a standing desk. Regardless of the mechanism, there is a possibility that broadly advocating for prolonged standing as an alternative to prolonged sitting may have several unintended consequences that negatively affect the health of the worker. These negative side effects may be more pronounced in female workers, and our data indicate that public health initiatives may need to be more cautious about advocating for standing as an alternative to sitting.

Despite possible differences in orthostatic tolerance, we did not observe any significant differences in vascular function between male and female participants. Previous research has indicated there is a lower prevalence of peripheral artery disease in females (Fowkes et al., 1991). The clinical and prognostic value of FMD and PWV for the approximation of cardiovascular risk has been determined in both health and disease (Ras et al., 2013). Indeed, endothelial dysfunction, FMD, tends to precede atherogenesis (Widlansky et al., 2003), and the leg vasculature is more susceptible to the development of atherosclerosis compared to other vessels (Padilla & Fadel, 2017). Previous research indicating sitting-induced impairments in FMD are reduced in females (Vranish et al., 2017) implies the cardiovascular impact of sedentary behavior differs between males and females. In contrast, our study indicates that sitting and standing induce similar responses in FMD and PWV respectively. Further, both the male and female participants equally benefitted from passive heating and standing in regard to FMD. However, how these effects relate to a sedentary lifestyle, wherein longer periods of sitting are repeated daily over weeks/months/years, needs to be explored.

### Considerations

There are three primary differences between our study and other studies investigating sitting induced vascular dysfunction: location of FMD measurement, duration of sitting, and our sample of female participants. In our study, FMD was assessed at the superficial femoral artery, but most studies have assessed lower limb FMD at the popliteal artery. We chose the SFA over the popliteal site based on current FMD guidelines indicating that the SFA accurately reflect endothelial function while the same has not been established at the popliteal site (Thijssen et al., 2011). If arterial bending causes endothelial dysfunction then it follows that greater dysfunction would be observed when FMD is measured at the popliteal artery where physical bending is occurring (i.e., at the knee). In addition, most studies subjected participants to longer durations of sitting (3-8h) than our 2-h protocol. This was not feasible in this protocol due to increased subject discomfort when they were asked to stand longer than 2 hours in our pilot work. The sitting and standing periods needed to be the same time period to make comparisons. Therefore, the lack of a statistically significant decline in FMD may be a result of differences in the arterial site being imaged via ultrasound, or a smaller effect due to the shorter duration of sitting. Further, previous work (Vranish et al., 2017) comparing the differences between the biological sexes recruited female participants who were eumenorrheic while our sample of participants were all taking a form of continuous hormonal contraceptive wherein they did not experience a regular menstrual cycle. It is possible that the differences in sex hormones between these studies contributed to the lack of differences in FMD between males and females in the current study.

### Conclusion

Standing and passive heating are equally effective at improving FMD in healthy young adults compared to prolonged sitting without passive heating. However, standing increases peripheral PWV, and increasing shear rate via passive heating does not attenuate this response. Meanwhile, our data indicate that circulating ET-1 is unaffected by either 2-h of sitting or standing. Overall, interventions for preventing sitting induced vascular dysfunction should focus on modifying shear rate (antegrade or retrograde) while preventing vasoconstriction which can temporarily increase lower limb arterial stiffness. These interventions could be in the form of local heating or standing (similar to this study), or may be as simple as physical activity breaks which can also increase antegrade shear rate and improve FMD. Contrary to previous work, we found the responses in FMD, PWV, and ET-1 were similar between sexes. Collectively, our findings indicate a lack of sex differences in vascular responses to 2-h of prolonged sitting or standing and provide novel information about the vascular consequences to this everyday hemodynamic challenge in young male and female adults. Further, the potential interventions, local heating and standing, resulted in similar outcomes for males and females. However, the experience of side effects may be more pronounced in females, and tailoring alternatives to sitting based on biological sex may be necessary in order to avoid the potential of syncope during prolonged standing.

## Additional Information

This article appears as a preprint: Caldwell, A.R., Jansen, L.T., Rosa-Caldwell, M.E., Howie, E.K., Gallagher, K.M., Turner, R.C., Ganio, M.S. Combined Effects of Standing and Passive Heating on Attenuating Sitting-Induced Vascular Dysfunction. bioRxiv. DOI: TBA

## Author Contributions

All authors meet the standards for authorship and all those who qualify are listed. All authors approved of the final version of this manuscript and agree to be accountable for all aspects of the work.

Contributed to conception and design: ARC, MSG, KMG, EKH, RCT

Contributed to acquisition of data: ARC, LTJ, MERC

Contributed to analysis and interpretation of data: ARC, RCT, MSG, KMG, EKH, MERC, LTJ

Drafted and/or revised the article: ARC, RCT, MSG, KMG, EKH, MERC, LTJ

## Funding

Partial funding for this study was provided by the Ph.D. student Research Award from the Central States Chapter of the American College of Sports Medicine.

### Acknowledgements

We would like to thank all the personnel and staff at the Exercise Science Research Center, and in particular, Dr. Michelle Gray and Shari Witherspoon for their support. We would also like to thank our wonderful undergraduate students Jeff Rogers, Zackary Vaughn, and Garrett Pierce for their help with data collection and the countless hours spent analyzing ultrasound videos.

